# Ensemble Classification through Random Projections for single-cell RNA-seq data

**DOI:** 10.1101/2020.06.24.169136

**Authors:** Aristidis G. Vrahatis, Sotiris Tasoulis, Spiros Georgakopoulos, Vassilis Plagianakos

**Affiliations:** Department of Computer Science and Biomedical Informatics, University of Thessaly, Lamia, Greece

**Keywords:** Classification, Visualization, Big Data, Big Biomedical Data, Transcriptomics, Single-cell RNA-seq, High Dimensional Data

## Abstract

Nowadays the biomedical data are generated exponentially, creating datasets for analysis with ultra-high dimensionality and complexity. This revolution, which has been caused by recent advents in biotechnologies, has driven to big-data and data-driven computational approaches. An indicative example is the emerging single-cell RNA-sequencing (scRNA-seq) technology, which isolates and measures individual cells. Although scRNA-seq has revolutionized the biotechnology domain, such data computational analysis is a major challenge because of their ultra-high dimensionality and complexity. Following this direction, in this work we study the properties, effectiveness and generalization of the recently proposed MRPV algorithm for single cell RNA-seq data. MRPV is an ensemble classification technique utilizing multiple ultra-low dimensional Random Projected spaces. A given classifier determines the class for each sample for all independent spaces while a majority voting scheme defines their predominant class. We show that Random Projection ensembles offer a platform not only for a low computational time analysis but also for enhancing classification performance. The developed methodologies were applied to four real biomedical high dimensional data from single-cell RNA-seq studies and compared against well-known and similar classification tools. Experimental results showed that based on simplistic tools we can create a computationally fast, simple, yet effective approach for single cell RNA-seq data with ultra-high dimensionality.

## 1 Introduction

Ongoing technological advances in the biomedical research field have increased the amount of the produced data to such an extent that research has shifted to data-driven computational methodologies. This evolution has revolutionized several areas including the biomedical domain [1] along with its important sub-disciplines such as bioinformatics, clinical informatics, imaging informatics, and public health informatics [2]. Regarding the bioinformatics aspect, expression profiles through omics studies (genomics, transcriptomics, proteomics, etc.) consist of an important part of the biomedical data expansion. The main reason is the Human Genome Project which changed the way we approach the interpretation of complex diseases and biological processes [3]. The technological development of recent decades on this field has created a large pool of ultra-high dimensional datasets. As the DNA sequencing cost significantly drops each year [4], the data volume increments relatively. Therefore the need arises to create new computational methods under the perspective of Machine Learning to uncover the data volume and complexity.

Nowadays, we are in the era of the emerging single-cell RNA-sequencing (scRNA-seq) technology, which belongs to the family of next-generation sequencing (NGS) technologies. he main difference with traditional methods, which export measurements from a bulk of cells, is that isolates individual cells offering gene expressions for each one. Through the scRNA-seq data we can achieve a deeper knowledge of the cellular functions opening new roads for the discovery of novel disease biomarkers. In the computation perspective, scRNA-seq studies generate a large amount of data with several cells and tens on thousands gene’s measurements. ost scRNA-seq data so far are belong to the “small n large p” category, where n is the number of samples (cells) and p is the number of dimensions (genes). ow the technological evolution allow us to manage datasets with with hundreds of thousands or even millions of cell samples. In both cases, we have datasets with ultra-high dimensionality, where there is a great risk of falling into the “curse of dimensionality” trap.

Classifying scRNA-seq data is a major computational challenge for this optimized NGS technology, while they explore the cellular differences offering in a higher resolution for the main building blocks of organisms. There is remarkable progress in classification methods of gene expressions data for single-cell RNA-sequencing studies [5, 6], however this field in its infancy.

In this work, we study the classification of single-cell RNA-seq data focusing in particular, on their high-dimensionality aspect. We argue that the recent MRPV (Multiple Random Projections with Voting) [7] classification scheme, has the true potential to be established as the new default in dealing with biomedical tasks with similar characteristics. MRPV main properties are its simplicity, its computational speed, its scalability, and its accuracy even on ultra-high dimensional data, rendering as a reliable and robust tool.

In what follows, Section 2 is providing details regarding the background material for MRPV. Next in section 3 we describe the MRPV algorithmic procedure in every detail while Section 4 is devoted to the experimental analysis and evaluation. Extensive commenting and discussion regarding the results is presented in Section5. The paper concludes in Section 6 with remarks and future directions.

## 2 Background Material: The Random Projection Method

Dimensionality reduction methods offer the potential to discover the structure of ultra-high dimensional data. The most common procedure for dimensionality reduction involves the projection of the data onto a lower dimensional orthogonal subspace. The Random Projection (RP) method [8–10], has been appeared to have the upside of great information portrayals with negligible calculation exertion, for information measurements in the scope of many thousands or even millions. In RP, the original high dimensional data is projected onto a lower dimensional subspace using a random matrix whose columns have unit lengths. The core idea behind it is given by the Johnson-Lindenstrauss lemma [11], which states that we can reduce the dimension dramatically and still be sure that the pairwise distances we can reduce the dimension by a number of orders of magnitude, while potentially almost preserving the statistical efficiency. More specifically, given a set of *n* points in a high dimensional Euclidean space with *a* dimensions, it can be mapped down onto an *r < O*(log *n/ε*^2^) dimensional space such that the distances are not distorted more than a factor of 1 *± ε*, for any 0 *< ε <* 1. It is interesting to note that the lower bound on *r* in the Johnson–Lindenstrauss lemma does not depend on *a*.

The RP method makes a perfect candidate when dealing with millions of dimensions since its inherent simplicity allows parallel implementation, a very crucial feature when we have to deal with computational complexity and memory management issues. Studies have been shown its similar behavior or superiority over the widely-used Principal Component Analysis (PCA). Indicatively, it has been shown that RP preserves distances having a comparable to PCA performance on image and text data operating with a much shorter execution time [9]. Furthermore, studies have been provided evidence that in a data clustering process, the clusters tend to appear a more spherical structure in the resulting dimension using the Random Projection method [11, 12]. These studies showed that RP performs better than PCA on eccentric data, where PCA might fail.

The main computational bottleneck of the RP is the orthonormalization of the random matrix *R*, however, for very large *a*, we can avoid it. As shown in [13], in high dimension spaces, there exists a much larger number of almost orthogonal vectors than orthogonal directions. Thus, high-dimensional vectors having random directions are very likely to be close to orthogonal. Regarding the elements of the matrix *R*, it has been shown that it is not necessary to be drawn from a Normal distribution. More efficient schemes have been proposed [8] imposing further computational benefits. Forming the random matrix *R* and projection the *n × a* data matrix *D* onto *r* dimensions is of order *O*(*arn*), and if the data matrix *D* is sparse with about *c* non zero entries per column, the complexity is reduced to *O*(*crn*) [10].

Random projections are combined with hierarchical clustering in [14] to offer expedient construction of dendrograms with provable quality guarantees. In [15] such random projections used to develop clustering algorithms that significantly improve run-time. It has also been shown that the random projection method can be applied before the *k*-means clustering with great success [16]. RP was coupled with a hierarchical data clustering algorithm in [17] showing that the theoretical properties of both components can be combined in a promising and effective clustering framework, orders of magnitude faster, while in [18] similar methodology was applied on ultra high dimensional data exposing its extreme parallelization and efficient memory management. In all aforementioned cases RP was utilized in unsupervised learning schemes. Since the random space is generated without considering the structure of original data may not capture task-related information, as such the benefits with respect to classification accuracy in using RP in supervised learning are not obvious. Nevertheless, a class of methods which involves the employment of RP in an ensemble scheme by applying the RP method multiple times on the same data aiming in achieving superior statistical accuracy, have recently emerged with very promising results.

### 2.1 Random Projections for Ensemble Classification

Random projections for the construction of an ensemble model have been used in [19] where the authors reported the reliability and efficiency of the concept, but without considering real high dimensional applications. Cannings et al. [20] proposed an ensemble methodology for high-dimensional data classification providing evidence that the straightforward aggregation is not always sensible. They claimed that possible bad projections can affect negatively the ensemble classifier, as such they proposed a methodology that minimizes the test error, closely related to the projection pursuit concept. In addition, instead of employing a simple majority vote for aggregation, the voting threshold is chosen in a data-driven fashion in an attempt to minimize the test error of the infinite-simulation version of the random projection ensemble classifier. As a result, although these restrictions allow the establishment of theoretical guarantees several considerations arise for very high dimensional tasks with respect to computational complexity. More importantly, it is not clear whether sub-selecting optimal random projections is actually a step backwards, reducing diversity [21], which may result in diminishing the true benefits of an ensemble classifier.

More recently, it has been experimentally shown [7] that utilizing random projections ensembles along with a simplistic majority voting scheme on the classification results from all independent data spaces, we can achieve improved classification accuracy when tested on very high dimensional biomedical data, while simultaneously minimizing computational complexity.

Motivated by these promising early results [7] in this paper, we focus on extensively testing the performance of this approach for the task of Single cell data classification which due to its basic characteristics seems like a best fit challenge allowing us to expose its usefulness in similar small “n” large “p” tasks. In our advantage, the utilization of a more simplistic approach allows extensive parallelization and efficient memory management, a crucial feature when we deal with big data with ultra-high dimensionality. Thus we describe the model and provide the implementation that incorporates such optimizations.

## 3 The Multiple Random Projection and Voting Methodology

The Multiple Random Projection and Voting (MRPV) framework consists of three main steps, i) the creation of multiple random subspaces, ii) the train step of a classifier in each space and iii) the test step for the new sample and the voting process to identify its dominant class for each sample. A flowchart diagram of the complete methodology is presented in Figure 1.

**Figure 1:**
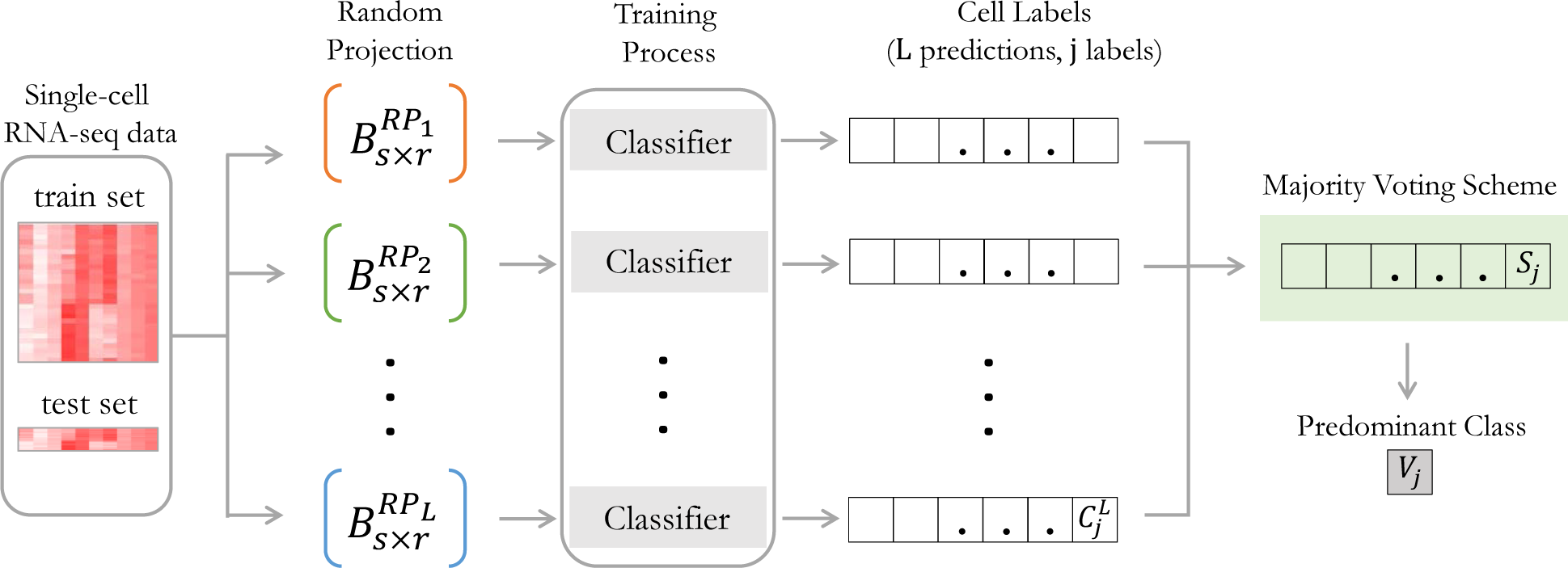
MRPV framework overview

Let *A ∈* ℝ^*n×d*^, be the original single-cell RNA-seq dataset with *n* samples in *d* dimensions. Let 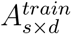 denote the *s* labeled samples data with *d* gene expressions used for training any classifier and 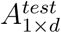 denote the query sample vector (which may also be a batch of samples constituting a matrix). The projections of 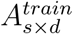 for all random subspaces can be calculated in a single matrix multiplication using a random frame *R*_*d*×*h*_, where *h* = *r* × *l* and *l* is the number of *r*-dimensional random projections (*r* ≪ *d*).

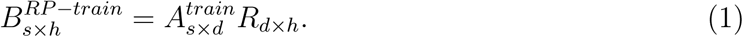

Then, we independently train the classifier for each projection by sub-selecting the corresponding columns of *B*^*RP*−*train*^. Subsequently, for each test point we perform a vector by matrix multiplication:

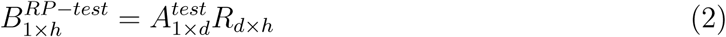

and then the appropriate columns of 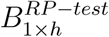 are selected for each corresponding classifier. Through this process we gain the advantage of utilizing optimized matrix operations libraries that significantly speeding up the procedure. In particular, for big datasets, when memory requirements cannot be fulfilled, it has been shown [18] that by diminishing and reconstructing the random matrix *R* using the random generator seed allows parallel execution in distributed environments.

The MRPV approach belongs to the “parallel ensemble methods” category for which the base classifiers are constructed in parallel exploiting independence. Similarly to bagging, we expect that by averaging (majority voting), the classification error will be reduced. We particularly focus on an ensemble methodology that uses a single base learning algorithm applied to the set of generated random projections. In our methodology, we generate several independent *d* × *r* random matrices and calculate the corresponding *n* × *r* projections for each one of them. Here, it is important to notice that we do not require any level of approximation of the pairwise distances in the projected space, thus the resulting dimensionality *r* is no longer bounded by *O*(log *n/ε*^2^), while *R* does not need to be orthonormal.

We experimentally show that maximum classification accuracy can be achieved even for very low *r* values, allowing us to dramatically reduce the computational complexity. As such, the dimensionality of each random projection *r* and their count *l* used in the scheme remain user-specified parameters, discussed further in Section 4.

### Algorithm 1 MRPV

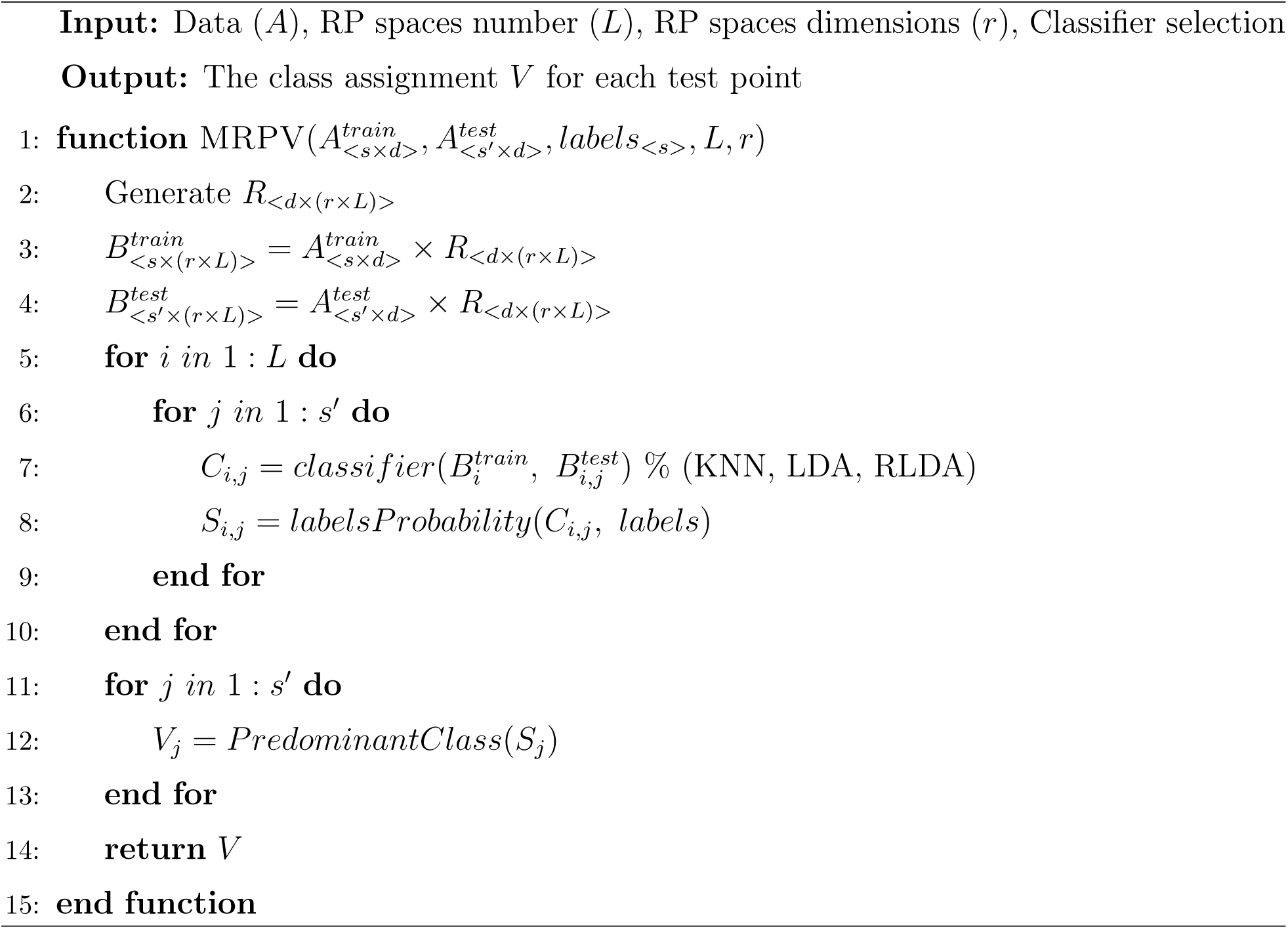

## 4 Experimental Analysis

### 4.1 Dataset Description

The performance of the proposed MRPV was evaluated on four real transcriptomics datasets from single-cell RNA-seq studies (Table 1). Datasets were obtained from Gene Expression Omnibus (GEO), the widely-used database repository for high throughput gene expression data.

**Table 1:** Attributes of transcriptomics experimental data

| GEO Dataset | Type | Dimensions | Samples | Classes | References |
| --- | --- | --- | --- | --- | --- |
| GSE103334 | scRNA-seq | 23,951 | 2,208 | 4 | [22] |
| GSE86469 | scRNA-seq | 26,616 | 638 | 2 | [23] |
| GSE59739 | scRNA-seq | 25,333 | 854 | 3 | [24] |
| GSE52583 | scRNA-seq | 23,228 | 201 | 4 | [25] |

**Table 2:**
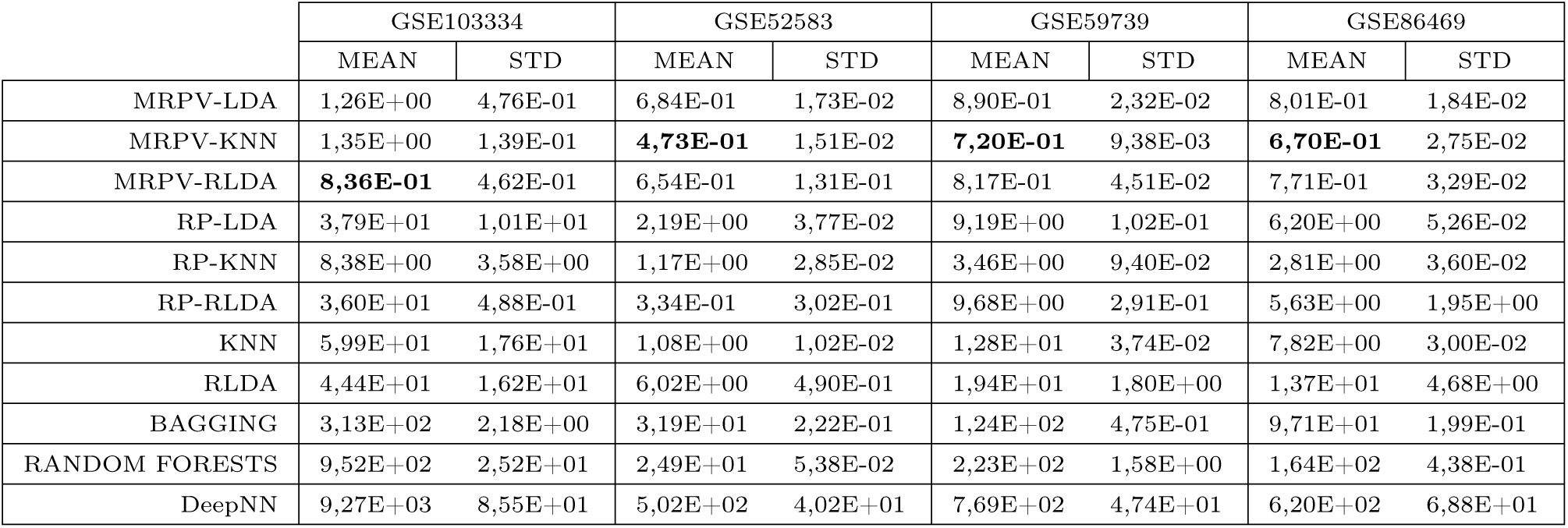
Mean and standard deviation (STD) values of the computational time performance of MRPV frameworks along with the other similar algorithms used as comparative evaluation in the current work. All algorithms have been executed 100 times independently for all four single-cell RNA-seq dataset. It is observed that our framework has the shortest execution times, especially the MRPV-KNN. his confirms the simplicity and the low computational effort required in random projections. Although in the RP-based algorithms one random projection is applied, our model is faster since based on our assumptions, we perform projections in extremely small dimensions, where the simple Random Projection method cannot cannot be guaranteed its reliability.

|  | GSE103334 |  | GSE52583 |  | GSE59739 |  | GSE86469 |  |
| --- | --- | --- | --- | --- | --- | --- | --- | --- |
|  | MEAN | STD | MEAN | STD | MEAN | STD | MEAN | STD |
| MRPV-LDA | 1,26E+00 | 4,76E-01 | 6,84E-01 | 1,73E-02 | 8,90E-01 | 2,32E-02 | 8,01E-01 | 1,84E-02 |
| MRPV-KNN | 1,35E+00 | 1,39E-01 | **4,73E-01** | 1,51E-02 | **7,20E-01** | 9,38E-03 | **6,70E-01** | 2,75E-02 |
| MRPV-RLDA | **8,36E-01** | 4,62E-01 | 6,54E-01 | 1,31E-01 | 8,17E-01 | 4,51E-02 | 7,71E-01 | 3,29E-02 |
| RP-LDA | 3,79E+01 | 1,01E+01 | 2,19E+00 | 3,77E-02 | 9,19E+00 | 1,02E-01 | 6,20E+00 | 5,26E-02 |
| RP-KNN | 8,38E+00 | 3,58E+00 | 1,17E+00 | 2,85E-02 | 3,46E+00 | 9,40E-02 | 2,81E+00 | 3,60E-02 |
| RP-RLDA | 3,60E+01 | 4,88E-01 | 3,34E-01 | 3,02E-01 | 9,68E+00 | 2,91E-01 | 5,63E+00 | 1,95E+00 |
| KNN | 5,99E+01 | 1,76E+01 | 1,08E+00 | 1,02E-02 | 1,28E+01 | 3,74E-02 | 7,82E+00 | 3,00E-02 |
| RLDA | 4,44E+01 | 1,62E+01 | 6,02E+00 | 4,90E-01 | 1,94E+01 | 1,80E+00 | 1,37E+01 | 4,68E+00 |
| BAGGING | 3,13E+02 | 2,18E+00 | 3,19E+01 | 2,22E-01 | 1,24E+02 | 4,75E-01 | 9,71E+01 | 1,99E-01 |
| RANDOM FORESTS | 9,52E+02 | 2,52E+01 | 2,49E+01 | 5,38E-02 | 2,23E+02 | 1,58E+00 | 1,64E+02 | 4,38E-01 |
| DeepNN | 9,27E+03 | 8,55E+01 | 5,02E+02 | 4,02E+01 | 7,69E+02 | 4,74E+01 | 6,20E+02 | 6,88E+01 |

More specific, the first dataset (accession number: GSE103334) has transcriptomics experimental data profiles for 23, 951 genes from 2, 208 cells separating in 4 different time periods [22]. Data came from time-points samples of Mus musculus (mmu) organism, during the progression of neurodegeneration, i) before p25 induction (0 weeks), ii) 1 week, iii) 2 weeks, and iv) 6 weeks after p25 induction. The first three categories had 576 cell samples and the latter 480. The main scope of the analysis of these expression profiles by throughput sequencing at single-cell level was the temporal record of microglia activation in neurodegeneration.

The single-cell RNA seq data with accession number GSE86469, includes gene expressions of 26, 616 genes for 638 cell samples. These are samples from human pancreatic islet obtained from non-diabetic (ND) and type 2 diabetic (T2D) cadaveric organ donors. Here the scope is to achieve the discrimination of ND and T2D cell types. The single-cell RNA seq data with accession numbers GSE59739 [24] and GSE52583 [25], contain single-cell RNA seq measurements for 25, 333 and 23, 228 genes respectively. The measurements are on 854 and 201 samples, separated at 3 and 4 classes respectively. Both data are from mouse models (Mus musculus) as well as coming from lumbar dorsal root ganglion and distal lung epithelium cells respectively.

We aim to show how the proposed RP-ensemble scheme affects the performance of other well-established classification tools in general, such as the kNN classification method. Also, since we have observed that minor changes for each parameter selection do not significantly alter the results, we chose to exclude an extensive analysis in an attempt to avoid further confusion upon the paper contributions.

### 4.2 MRPV Framework Performance

The performance of the MRPV framework was evaluated against eight well-established and similar classification methods. he rationale behind the optimal selection of algorithms was to cover the entire range of various categorization types as well as to have a fair comparison with similar tools. We select the three main classifiers of our framework (LDA, KNN, RLDA) using the simple Random Projection method, a deep learning, a bootstrap aggregating (bagging) and a tree-based classifier. We also include the traditional KNN and Regularized linear discriminant analysis (RLDA) classifiers, excluding the traditional LDA classifier since its execution requires ultra-high amount of computational resources and it is too far beyond a compatible processor. Overall, the comparisons were made with the (a) LDA with the Random Projection method (RP-LDA), (b) KNN with the Random Projection method (RP-kNN), (c) RLDA with the Random Projection method (RP-RLDA), (d) KNN, (e) RLDA, (f) BAGGING, (g) RANDOM-FORESTS, and (h) Deep Neural Network (DeepNN).

Following, we provide a brief description of each method along with their parameter selection. The KNN performs k-nearest-neighbor classification model [26] using the default parameters with Euclidean as distance measure as well kdtree option as search method for *N* = 5 nearest neighbors. The RLDA performs the regularized LDA, where all classes have the same covariance matrix 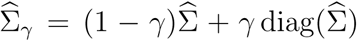, with 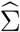 be the empirical, pooled covariance matrix and is the amount of regularization.

RP-LDA, RP-KNN and RP-RLDA perform LDA, KNN, and RLDA respectively on one projected space by applying the standard random projection method. The size of the resulting subspace *r* is defined by r = 4 (*ε*^2^*/*2 − *ε*^3^*/*3)^−1^ ln(n) [27]. The variable *n* indicates the sample size while the parameter *ε* was set *ε* = 0.2 in order to achieve a significant reduction in dimensionality that would allow faster execution times.

Deep Neural Networks are utilized with four neural layers of 100 neurons each one due to the high-dimensionality of the datasets. DNN was applied with hyperbolic tangent activation functions for each layer while the softmax functions utilized on the output layer. The training of the model was based on ADAM [28] learning algorithm using the value 0.001 for both *β*_1_ and *β*_2_ learning rate parameters.

The bagging approach performs an ensemble learning method for classification using a bagging approach which stands for bootstrap aggregation [29]. The minimal leaf sizes for bagged trees was set to 1 and the number of predictors selected at random for every decision split with size equal to the square root of the number of features (genes). Random Forests perform an ensemble learning method for classification by constructing a multitude of decision trees at training time and outputting the class that is the mode of the classes (classification) of the individual trees [30]. Random Forests classification model is applied using 100 trees and the parameter m-try set to square root of the number of features (genes). Performance was assessed by measuring three indicative classification measures such as accuracy, specificity and F1-score, three well-known diagnostic tests for multi-class evaluation. Through these measures we have a clear view about the strengths and weaknesses of each method. All executions were made using the 10-fold cross validation process in 100 independent trials. All algorithms were run with the corresponding optimal parameters. All algorithms, besides MRPV-LDA, MRPV-KNN and MRPV-RLDA, were rerun with different parameters, though no significant improvement was detected. Boxplots depict the comparative results for all methods in all four datasets.

We employed the Wilcoxon signed-rank test (P-value < 0.05) after observing that the median differences between pairs of observations (between MRPV frameworks and each algorithm under investigation) follow non-normal distribution (one-sample Kolmogorov–Smirnov test with P-value < 0.05). MRPV framework (MRPV-LDA, MRPV-KNN or MRPV-RLDA) performed better in almost all cases and the difference was statistically significant in most cases (see section 5).

## 5 Results and Discussion

We observe that both variations of MRPV are competitive against other methods for all datasets while all the three MRPV-based algorithms suggest an improvement over the straightforward application of LDA, KNN, and RLDA respectively. We interestingly observe that the proposed ensemble scheme suggests a more valuable integration of Random Projections than that of the basic RP method.

More specifically, for all three measures (Accuracy, Specificity, and F1-score), the MRPV-KNN had the best performance on GSE103334 with MRPV-LDA and MRPV-RLDA overcoming their relevant methods (RP-LDA and RP-RLDA respectively) in almost all cases. In all other datasets, the MRPV-RLDA outperforms all comparative methods with MRPV-LDA and MRPV-RLDA having similar behavior as previously. In few cases, we identify differences with no statistical significance (one-sample Kolmogorov–Smirnov test with P-value < 0.05) in three of the four datasets, such as (a) MRPV-LDA vs RP-LDA F1-score, MRPV-RLDA vs RP-RLDA F1-score on GSE103334, (ii) MRPV-KNN vs RP-KNN Accuracy and MRPV-KNN vs KNN accuracy, specificity values for all MRPV frameworks vs RP-based frameworks on GSE86469, (iii) MRPV-LDA vs RP-LDA Accuracy, MRPV-KNN vs RP-KNN Accuracy, MRPV-LDA vs RP-LDA specificity and F1-score on GSE52583 dataset.

Note here, that MRPV framework creates projected spaces with only 50 dimensions where the original datasets have tens of thousands of dimensions confirming the original hypothesis that high performance can be achieved even when significantly reducing the original dimensionality.

When comparing to the performance of the three well-established classifiers (Bagging, Random Forests, and Deep Neural Network) coming from diverse backgrounds, we conclude that the proposed approach still performs better enhancing its reliability. The stability of the proposed methods is also a matter of interest when comparing to the other methods that present a wide dispersion.

It is worth mentioning that the cell samples from all studies are not sufficiently separable and it is difficult to classify each cell category. This is illustrated quite clearly through the 2D visualization for the four single-cell RNA-seq datasets using well-known and cutting-edge visualization tools, in Figure 6. We selected PCA [31] and t-SNE [32], two well-known dimensionality reduction tools, UMAP [33] a state-of-the-art technique for dimension reduction and RPEV-MDS [34] a novel in-house 2D visualization tool with remarkable efficiency. s shown, class separation is not straightforward (especially for PCA) exposing the data complexity. The high intermix of classes across all visualizations confirms the data complexity and subsequently the challenging classification task.

**Figure 2:**
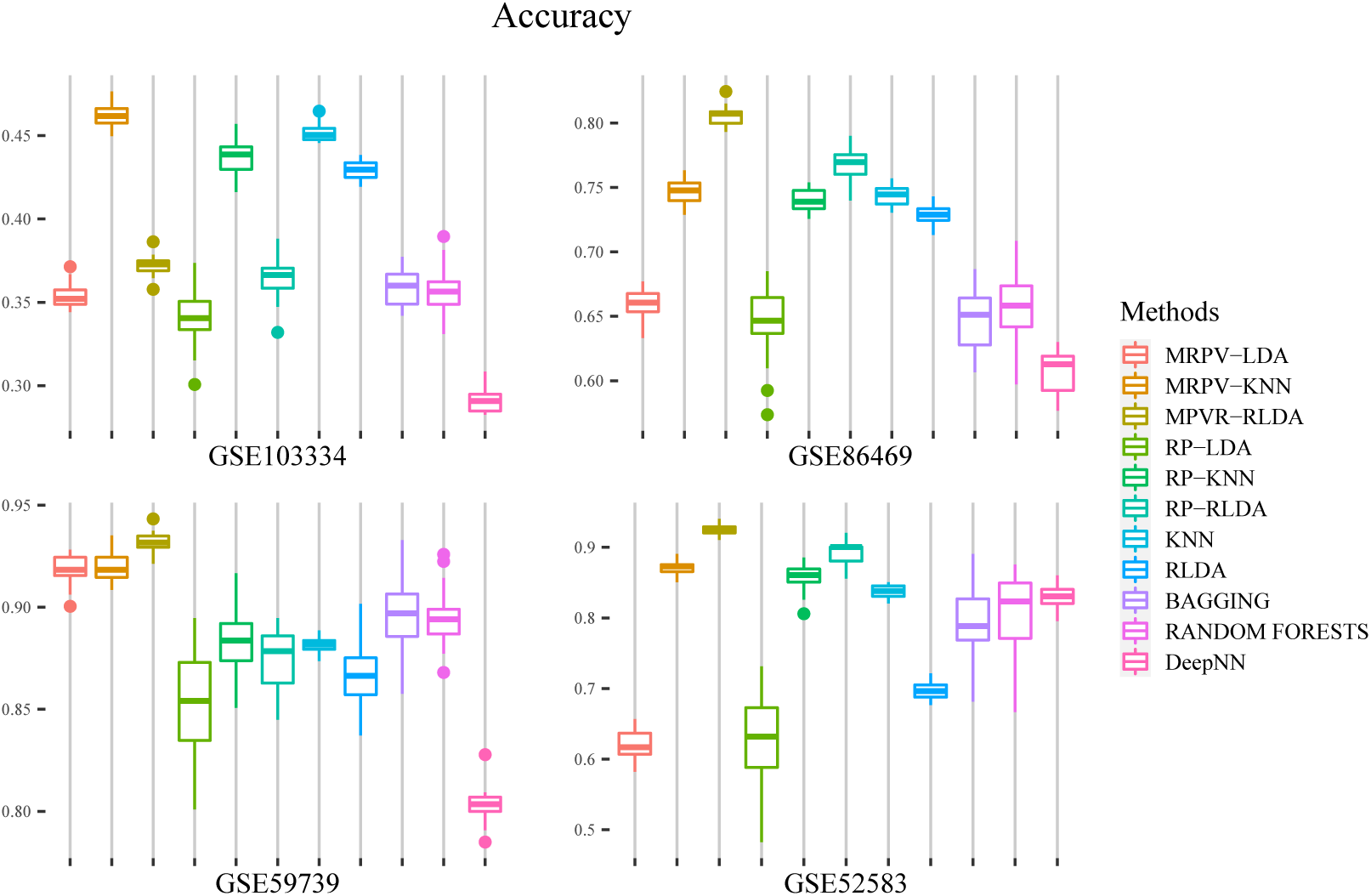
Classification performance of the proposed MRPV framework (MRPV-LDA, MRPV-KNN, MRPV-RLDA) compared to other eight classification methods for the dataset GSE103334, GSE86469, GSE59739 and GSE52583. Evaluation testing is made by examining Accuracy. Boxplots imprint the values via 100 independent executions for each dataset.

**Figure 3:**
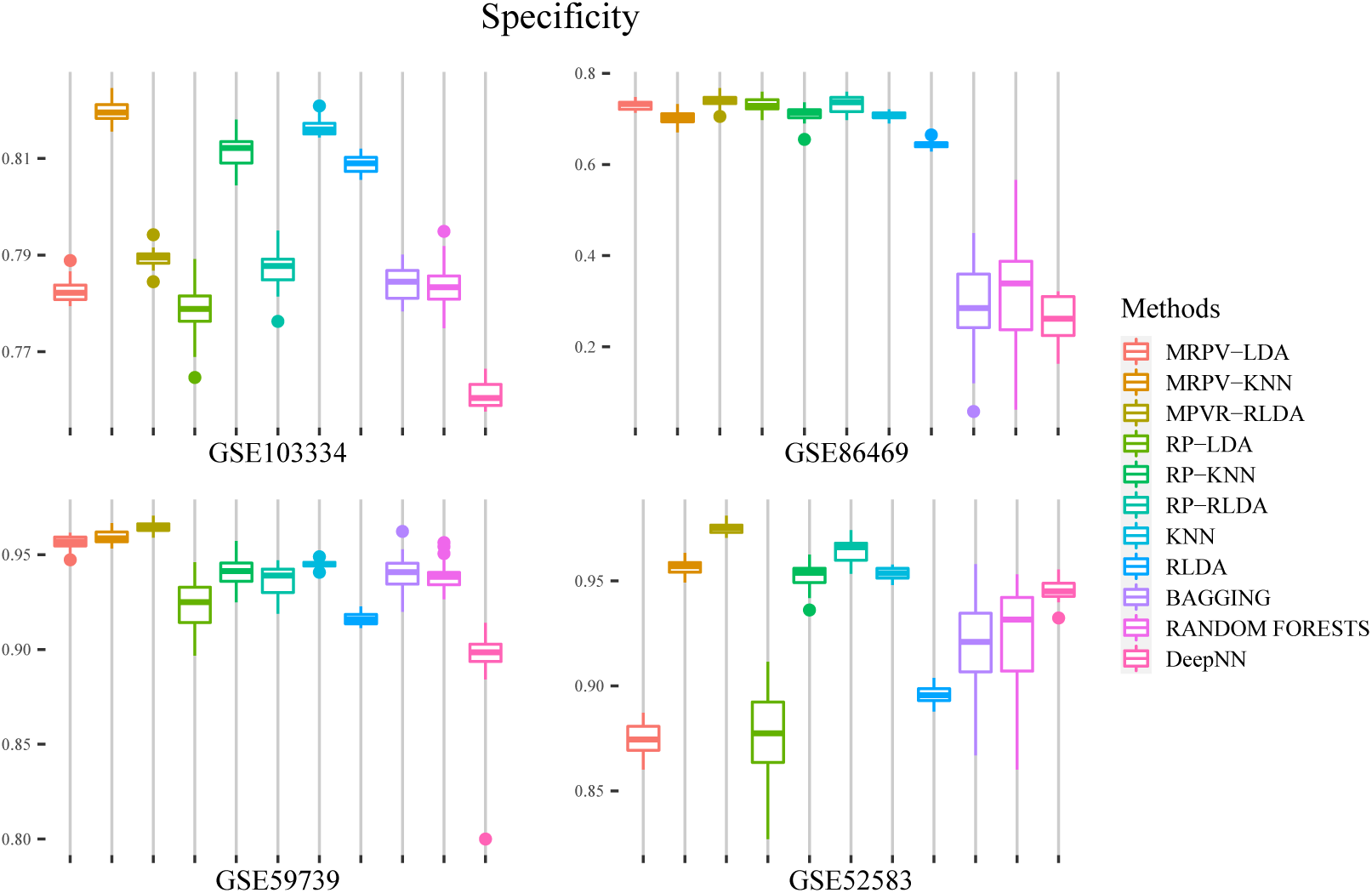
Classification performance of the proposed MRPV framework (MRPV-LDA, MRPV-KNN, MRPV-RLDA) compared to other eight classification methods for the dataset GSE103334, GSE86469, GSE59739 and GSE52583. Evaluation testing is made by examining Specificity. Box-plots imprint the values via 100 independent executions for each dataset.

**Figure 4:**
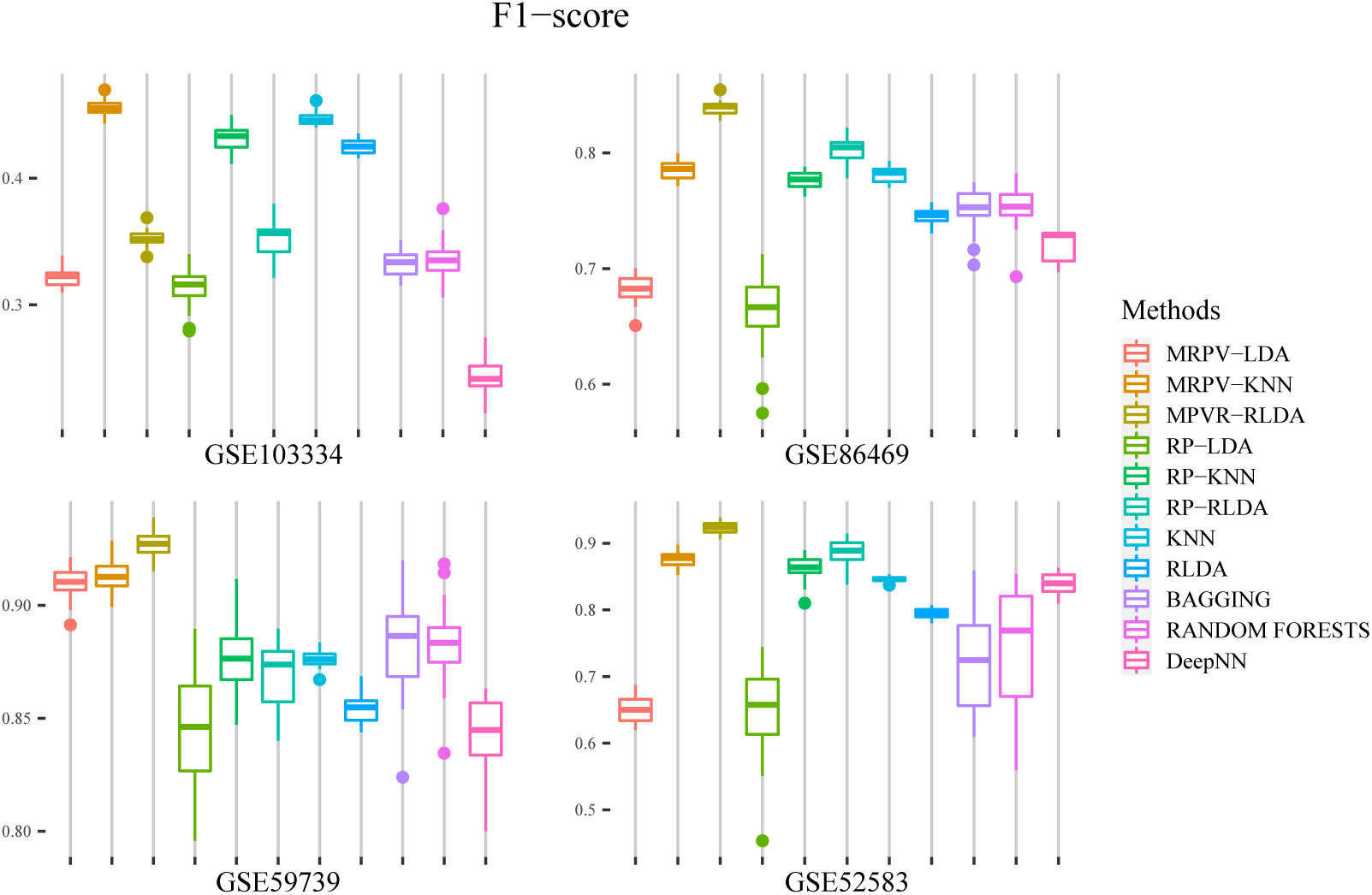
Classification performance of the proposed MRPV framework (MRPV-LDA, MRPV-KNN, MRPV-RLDA) compared to other eight classification methods for the dataset GSE103334, GSE86469, GSE59739 and GSE52583. Evaluation testing is made by examining F1-score. Boxplots imprint the values via 100 independent executions for each dataset.

**Figure 5:**
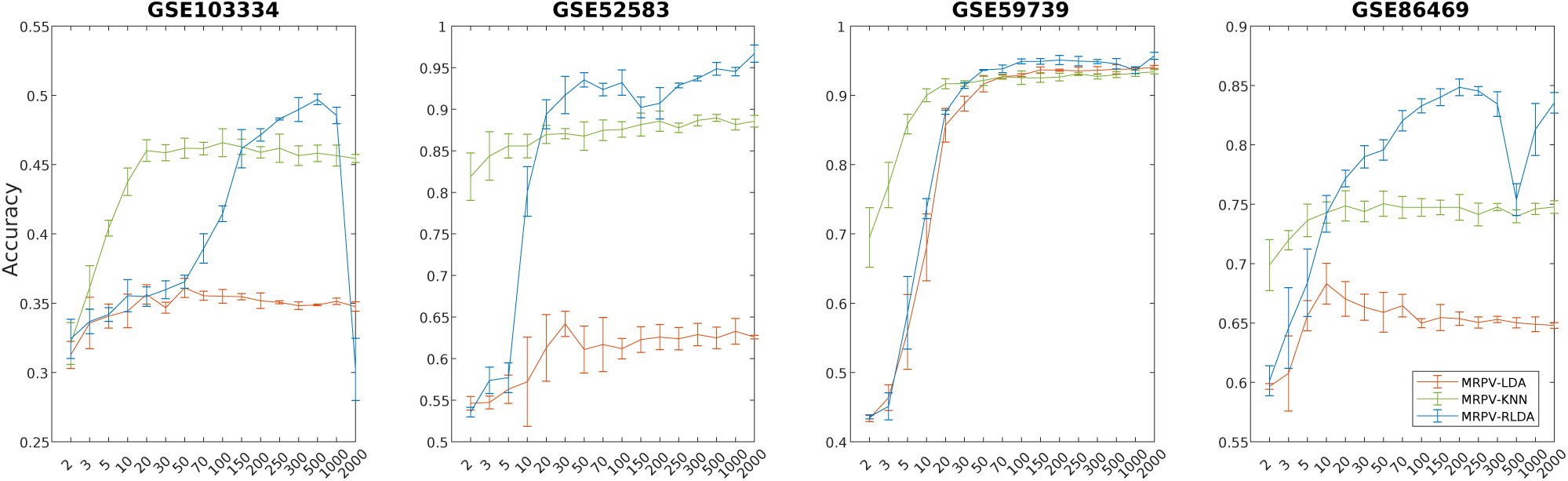
Error bars with the performance evaluation of the three MRPV-based algorithms accuracy (y axis) in terms of the dimensions number selection (x axis) of the random projection spaces. Starting from a quite small number of dimensions, we notice that the accuracy performance increases as the dimensions increase and after one point (about 30 to 50 dimensions), the accuracy performance stabilizes. So it can be concluded that MRPV model works well in the small dimensions without having to project the initial data to higher dimensions

**Figure 6:**
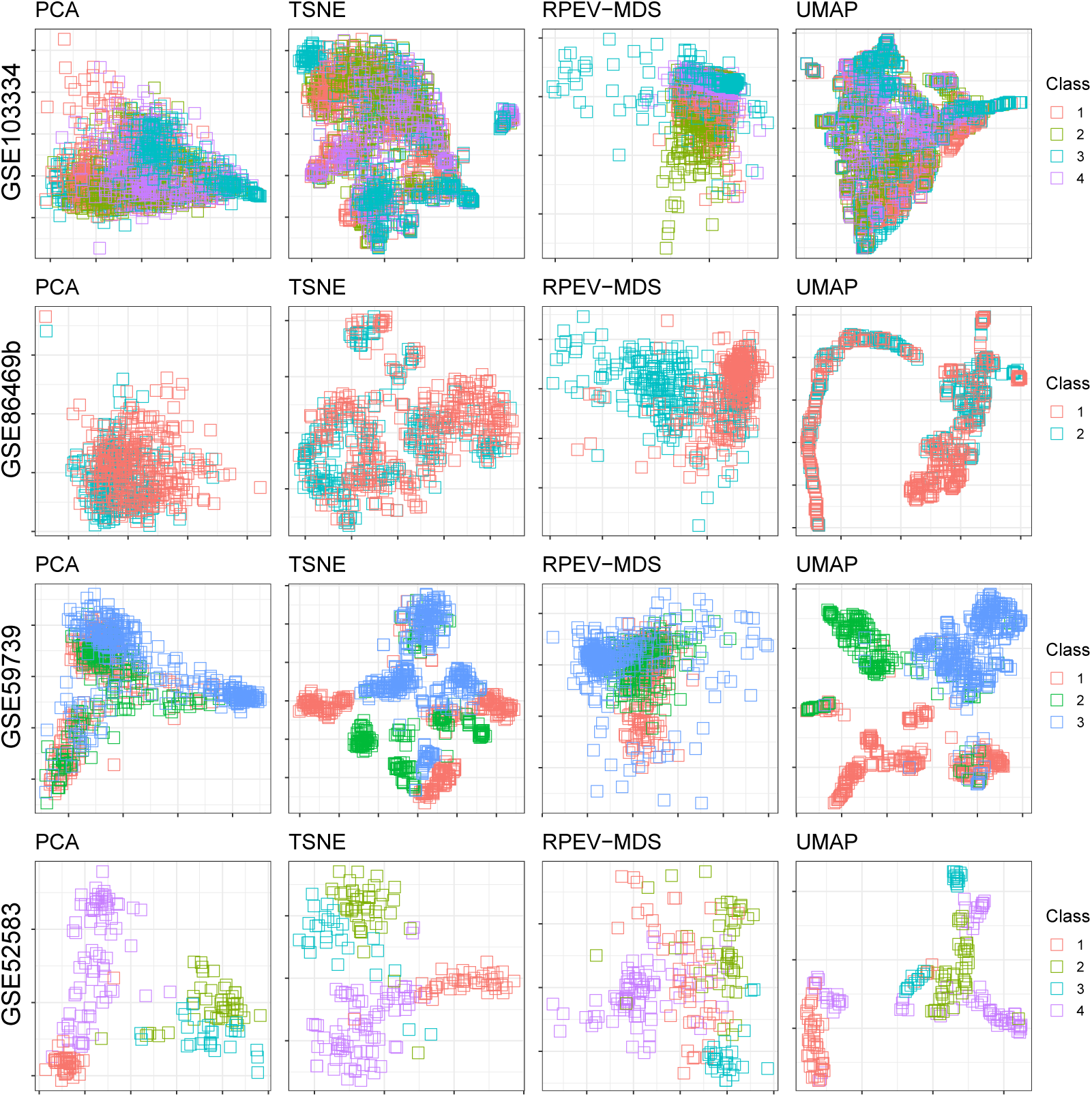
2D visualization for the four single-cell RNA-seq datasets using PCA, t-SNE, RPEV-MDS and UMAP tools. Each point represents a sample and each colour represents a different class type according to original data biologically meaningful annotation. We notice that the classes are not sufficiently separable, thus it is difficult to classify each cell category.

Considering the aforementioned results, we can clearly state that the proposed framework offers a promising approach in capturing the structure of the high-dimensional data maintaining its properties, while also suggesting great potential in minimizing computational complexity for many different classification approaches. he results show that MRPV is a reliable framework that can work efficiently in cases with significantly bigger data in terms of the number of samples is of much interest (small n – large p). In this case the dimensionality (number of genes here) can be much higher than the number of samples and we try to tackle problems related to the “curse of dimensionality”.

Note here, that MRPV framework allows the application of classification methods in ultra-high dimensional data which otherwise would be forbidden (mainly due to memory limitations). For two particular methods we show that simultaneously, improved classification accuracy can be achieved, indicating the potential of such approaches. In addition, the computational savings of the RP method are already justified in earlier works [35] along with our previous study [7] which we extend in this work. Through this study, we showed the superiority of MRPV framework compared to eight classification tools on four single-cell RNA-seq data.

Furthermore, to best of our knowledge, ensemble classification schemes using random projections have never been applied to big data with ultra-high dimensionality. The most indicative recent approach is the work of Cannings & Samworth [35] for which we provided several considerations arising for real world problems of ultra high dimensionality in terms of computational cost [7].

## 6 Conclusion

We presented the MRPV framework for the classification of big single-cell RNA-seq data with ultra-high dimensionality. Both are based on multiple random projections in a lower-dimensional space and on ensemble approaches that aim to exploit the different data structures obtained by each random projected space. The performance of our frameworks was evaluated in four public real experimental high-throughput biomedical data with transcriptomics expression profiles. The classification results showed the superiority of MRPV-KNN, MRPV-LDA and MRPV-RLDA against eight well-known tools. Most importantly, the proposed framework imposes further theoretical developments but also practical solutions for classification that can take advantage of its parallel fashion.

## Author Contributions

A.G.V. conceived of the study, designed the classification methodological framework, implemented the experimental analysis of the classification framework and drafted the manuscript. S.K.T. designed and implemented the visualization methodological framework, contributed in the interpretation of the results and the manuscript draft. S.V.G. contributed in the design and implementation of figures and contributed in the interpretation of the results. All the above actions were supervised by V.P.P. All authors read and approved the final manuscript.

## Acknowledgement

This research has been financially supported by the National Strategic Reference Framework (NSRF) Program with title: “Researcher Support with Emphasis on New Researches”, cofinanced by Greece and the European Union - European Social Fund.

